# An algorithm for predicting per-cell proteomic properties

**DOI:** 10.1101/2024.12.03.626698

**Authors:** José Ignacio Arroyo, Chris Kempes

## Abstract

Proteomic studies have traditionally focused on population-level analyses with an emphasis on the relative abundance of various proteins. Such studies have been useful in uncovering physiological differences across diverse species, the physiological response of individual species to distinct environmental conditions, and the function of individual proteins in the context of cellular networks. However, the absolute value of protein abundance in a cell is important for understanding single-cell physiology, detailed biophysical considerations, connecting diverse quantitative data, and for comparisons across species. Such detailed quantification will naturally occur as singlecell proteomics becomes more prevalent, but it is also of current interest to leverage population studies. There are several challenges here. First, most population studies do not measure the quantity of cells associated with a proteome. Second, recent work has shown that cell physiology radically shifts with cell size, and these effects need to be accounted for in going from population to single-cell estimates. Here we develop and implement a method to estimate the basic properties of proteomes, based on well-established scaling relationships among cell components, including genome size, cell size, and proteome volume. Our method estimates similar but higher total proteins per cell compared to previous theoretical and empirical estimations. Our algorithm has applications for interpreting proteomes, analyzing environmental samples, and designing artificial cells. While focusing on prokaryotes, we discuss how the method can be extended to unicellular and multicellular eukaryotes.

## Introduction

Proteomes are the total set of proteins in a cell, or populations of cells, in a given environment. Most proteomic studies quantify the proteome of a sample, e.g. a culture of bacterial cells or a sample of tissue. In this context, most proteomic studies quantify the total number of peptides in that sample and the relative abundance (i.e. the abundance of each protein divided by the total abundance) or a similar measure of each protein, such as proteins per million [1]. However, there are variety of biophysical, comparative, single-cell, and quantitative perspectives that require knowing the absolute abundance of proteins within individual cells [2–4]. For population studies, this means estimating the average abundance per cell, which can be calculated using the total population protein abundances and the population-level number of cells. Such a calculation is simple except that most studies do not report the number of cells [1, 2], except for recent single-cell proteomics studies that have thus far only been applied to *E. coli* [5]. Previous efforts have sought to quantify cell-level proteomics using the estimated number of molecules per cell volume [2] and here we expand on those perspectives using recently developed theory for bacteria. Specifically, we use well-established scaling relationships between the different components of the cell (DNA, proteome) and cell volume (e.g. [2–4, 6–9]), to scale the total abundance of proteins, the abundance of each protein, and the number of different proteins per cell volume.

Scaling laws are compact relationships for how features change with one another based on the size of the system. The simplest version of a scaling relationship is a power law which [10, 11] describes a pattern of the form

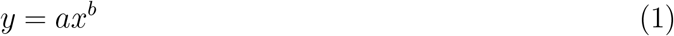

Where *y* is some cellular property, *a* is a normalization parameter, *x* is cell size, and *b* is a scaling exponent. Examples include the relationships between genome (amount of DNA) size and cell size, protein volume and cell size[3], or among RNA and proteins [8].

The exponent *b* accounts for the rate of change of *y* with *x*. For example, if *x* doubles in size and if *b <* 1, then *y* will increase by less than a factor of 2, whereas if *b* = 1 it will double, and if *b >* 1 it will increase by more than a factor of 2. Eq. 1 is often written logarithmically as *log*(*y*) = *log*(*a*) + *log*(*x*), and the parameters are fitted using simple linear regression [10]. There are two relevant properties of Eq. 1.; scale-invariance and universality. Scale invariance or self-similarity refers to the property of a function *f* (*x*) that under re-scalings of the variable *x* by a factor *λ* the function does not change, that is *f* (*λx*) = *λ*^*a*^*f* (*x*). Universality, often refers to the property of different systems having similar exponents. For instance, as described above, systems with sublinear exponents describe economies of scale if, for example, *y* represents a cost variable that does not grow proportionally with size. Often, scaling relationships that emerge from relevant principles that conform to complex systems such as optimization [9, 10, 12, 13].

Because these scaling relationships systematically interconnect many cellular features through the size of the system it is possible to estimate unmeasured features. Below we provide a method specifically aimed at estimating per-cell proteome properties. This method relies on inherent shifts in the protein composition with cell size. Specifically, as one goes from small species like *Mycoplasma* to larger species like *E. coli* - which traverses up to three orders of magnitude in cell volume - the protein concentration of cells radically dilutes [2, 3]. This is accompanied by shifts in all of the major macromolecules [3] and can be explained by certain considerations of optimal metabolism and physiology which change with cell size [3]. These relationships set a baseline for cellular concentrations and can be used to estimate the proteome of each cell. This approach will inherently be accompanied by some amount of error, and future studies should make concurrent measurements of features of interest.

## Results

### Mathematical framework

Our goal is to provide a simple algorithm for approximating the number of proteins in a cell, the abundance of each protein, and the maximum number of expressed proteins. Our approach is to use a combination of simple assumptions about the sample, and expectations from cellular scaling to estimate the proteome properties of a single cell. We begin by specifying the scaling relationships that allow us to interconvert between cellular properties and predict expectations for cells of any size, and then we will describe the algorithm that we developed to use these relationships to predict per-cell proteome quantities.

#### Total protein abundance

To calculate the number of proteins molecules in a cell we first calculate the cell volume from genome size, a quantity that is easily obtained from different databases (NCBI, for example). Then we use the relationship between protein volume and cell size, to calculate the predicted protein volume in a cell.

From genome size, *G*, (in Megabases; Mb), we estimate cell volume, *C*_*v*_ (*µ*m^3^), [14] as,

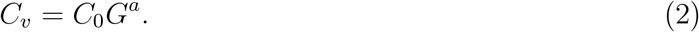

For prokaryotes, the parameter values in Eq. 2 are *a* = 3.52 and *C*_0_ = *e*^20.4^(1*/*978)^3.52^ (Table 1). Then, from cell volume the estimated total protein volume [3] is

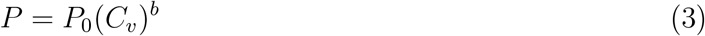

where *C*_*v*_ (*µ*m^3^) and P (m^3^). For prokaryotes the parameter values are *b* = 0.7 and *P*_0_ = 3.42*×*10^−7^(10^−18^)^0.7^ (Table 2).

**Table 1.** Variables, parameters, and constants of the mathematical framework for predicting “per cell” proteome properties.

| Quantity | Symbol | Unit |  |  |
| --- | --- | --- | --- | --- |
| **Variables** |  |  |  |  |
| Genome size | $G$ | Mbp | | |
| Cell volume | $C_v$ | $\mu\text{m}^3$ | | |
| Proteome volume | $P$ | $\text{m}^3$ | | |
| Vector of per-cell protein abund. | $\mathbf{N}$ | $\frac{\text{prot. molecules}}{\text{cell}}$ | | |
| Vector of obs. abundance | $\mathbf{M}$ | prot. copies | | |
| Total prot. abund. in the sample | $N_{sam}$ | $\frac{\text{prot.}}{\text{sample}}$ | | |
| Num. of prot. in the cell | $N_{cell}$ | prot. molecules | | |
| Num. of encoded prot.-coding genes | $P_{en}$ | $\frac{\text{enc. prot.}}{\text{genome}}$ | | |
| Num. of protein mol. in the sample | $P_{ex}$ | $\frac{\text{proteins}}{\text{sample}}$ | | |
| Coverage($=\frac{P_{ex}}{P_{en}}$) | $C$ | dimensionless | | |
| N. of diff. prot. in the sample | $D$ | $\frac{\text{prot. types}}{\text{Sample}}$ | | |
| N. of diff. obs. prot. in the sample | $D_{obs}$ | $\frac{\text{prot. types}}{\text{Sample}}$ | | |
| Max. number of dif. prot. per cell | $D'$ | $\frac{\text{prot. types}}{\text{cell}}$ | | |
| Correction factor | $\lambda$ | dimensionless | | |
| Residuals($= D_{obs} - D$) | $r$ | $\frac{\text{prot. types}}{\text{cell}}$ | | |
| **Parameters** |  |  | Value | Ref |
| Prefactor in the rel. $C_v \propto G$ | $C_0$ | $\frac{\mu\text{m}^3}{\text{Mbp}}$ | $2.15 \times 10^{-2}$ | [14] |
| Exponent in the rel. $C_v \propto G$ | $a$ | — | 3.52 | [14] |
| Prefactor in the rel. $P \propto C_v$ | $P_0$ | $\frac{\text{m}^3}{\mu\text{m}^3}$ | $8.59 \times 10^{-20}$ | [3] |
| Exponent in the rel. $P \propto C_v$ | $b$ | — | 0.7 | [3] |
| Ave. protein vol. for prok. | $P_{prok}$ | $\text{m}^3$ | $3.75 \times 10^{-26}$ | [15–17] |
| Prefactor in the rel. $P \propto G$ | $\delta$ | $\frac{\text{encoded proteins}}{\text{Mbp}}$ | 954.99 | TS |
| Exponent in the rel. $P \propto G$ | $\gamma$ | — | 0.96 | TS |
| Prefactor in the rel. $D \propto N_{cell}$ | $\alpha(C)_{C<0.5}$ | $\frac{\text{protein types}}{\text{protein molecules}}$ | — | — |
| Exponent in the rel. $D \propto N_{cell}$ | $\beta$ | — | — | TS |
| $\alpha$ for $C > 0.05$ | $\alpha_{C>0.5}$ | — | 9.33 | TS |
| Slo. of lin. the rel. between $\alpha \propto C$ | $m$ | $\frac{\text{protein types}}{\text{protein molecules}}$ | 12.95 | TS |
| Int. of the lin. rel. between $\alpha \propto C$ | $n$ | $\frac{\text{protein types}}{\text{protein molecules}}$ | -1.17 | TS |
| **Constants** |  |  |  |  |
| Average volume of an amino acid | $A_v$ | $\text{m}^3$ | $13.92 \times 10^{-28}$ | [15, 16] |
| Abbreviations; Num.:number, ind.:individual, prot.: proteins, abund.: abundance, diff.; different, TS: this study |  |  |  |  |

**Table 2.**
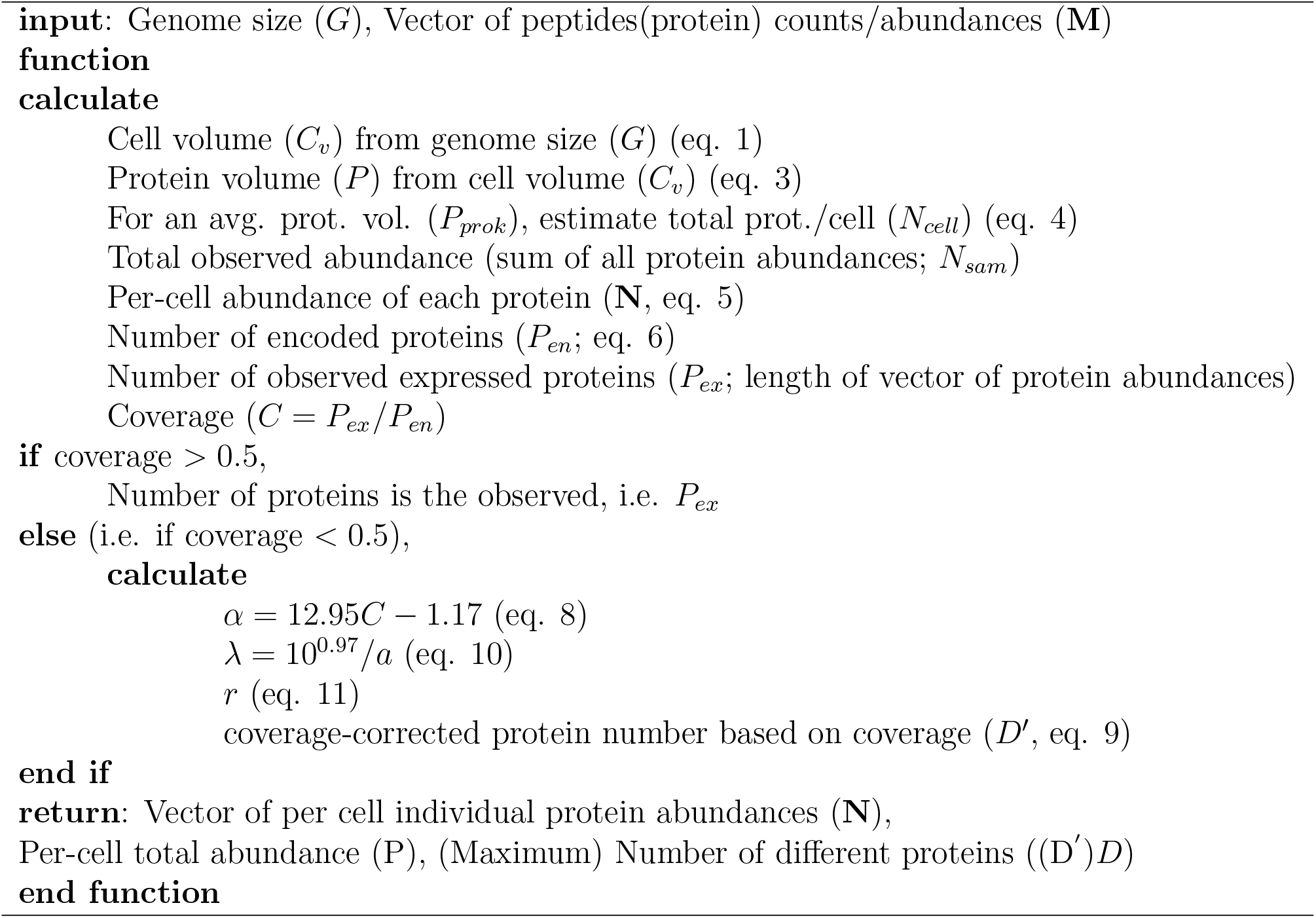
Algorithm/Pseudocode for proteomic data scaling.

#### Individual protein abundance

By multiplying the average volume of an amino acid by the average length (number of amino acids) of a protein, we calculate the average volume of a single protein. The average volume of an amino acid is 1.39×10^−28^*m*^3^ [15, 16], which mulpitplied by the average bacteria protein length, ≈270 [17], gives an average protein volume of 3.75 ×0^−26^*m*^3^. Dividing the total protein volume of a cell by the typical volume of an individual protein, the number of protein molecules is given by

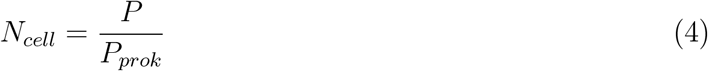

We will derive a formula to calculate the per-cell individual protein abundance. The observed abundance of each protein type is *m*_*i*_ and the predicted abundance of each protein type is *n*_*i*_, and as explained above the number of predicted proteins is *N*_*cell*_ and the number of proteins. We assume a direct proportion between the relative observed abundances and the relative predicted abundances, 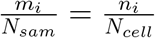. Then to calculate the predicted abundance of a protein type we simply multiply the observed abundance by the ratio of predicted to observed total abundances;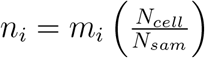.

As a single operation, we estimate the per-cell abundance of each of the proteins by multiplying the vector of the observed abundances of each protein in the sample (**M**) by the ratio of predicted proteins in the cell (*N*_*cell*_) and observed total proteins in the sample (*N*_*sam*_),

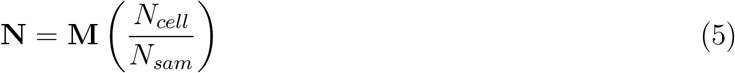

The resulting vector **N**, is the vector of abundances per-cell.

#### Number of different protein types

Often not every protein type (i.e. a given protein such as Fur or LysR) encoded in the genome is found in proteomes, either because some proteins are not expressed in a given environment or because of experimental limitations. For example, the amount of sample, and hence the number of cells in the sample, could affect not only the total abundance but the number of different expressed proteins observed. Consider a sample of the same species with a population of 100 cells versus a sample with 10,000,000. Likely, the sample of only 100 cells will not capture all the different types of proteins, especially low abundance ones, that a particular species can express under a given condition.

Most proteomic studies do not report the number of cells in the sample so we are not able to normalize the samples by number of cells, nor to quantify the associated error. The proportion of expressed proteins, *P*_*ex*_, over the number of encoded proteins, *P*_*en*_, is often known as the coverage, *C* = *P*_*ex*_*/P*_*en*_. In terms of data, *P*_*ex*_ corresponds to the length of the vector of protein abundances, and the value of *P*_*en*_ is obtained from known annotations/predictions made for genomes in databases such as the NBCI. We address these challenges by estimating *P*_*en*_ using the scaling between protein-coding genes and genome size, which is given by

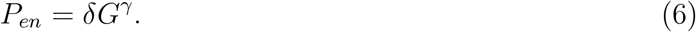

Although this relationship has been reported in the literature [18], the parameters *δ* and *γ* have not been estimated. We compiled data from representative prokaryotes (Supplementary Material) and found that *γ* = 0.96 and *δ* = 10^2.98^ (Table 1). Below we will show that this relationship is useful as just from genome size, we predict most of the per-cell quantities of proteomes.

To predict the number of different expressed proteins, *D*, and the total number of proteins, *N*_*cell*_, we empirically assessed the scaling relationship,

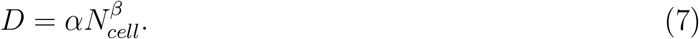

We found that there was a generally positive trend between the number of expressed proteins and total proteins but with a high variance. We observed that samples with low coverage were located in the lower region of this trend and samples with higher coverage were located in the upper region of this trend. To analyze this pattern more in detail, then we examined the relationship for different interval coverages of a size of 0.1, ranging from 0-0.1 to 0.8-0.9. We found that the prefactor varied with coverage, a relationship was linear up to a threshold of 0.5, after which the prefactor value remained mostly constant (SM Figure 1). On the other hand, the exponent remained constant with an average of 0.38 (SM Figure 1).

Accordingly, we re-express the relationship in Eq. (7) for data having a coverage above 0.5 (*C >* 0.5) and below 0.5 (*C <* 0.5) as,

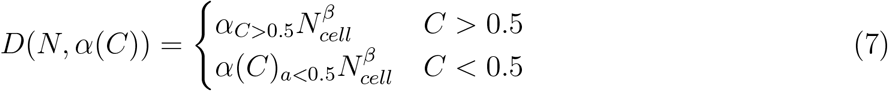

Here *α*_*C>*0.5_, and *β* stands for the prefactor and exponents for coverages greater than 0.5, and *α*(*C*)_*C<*0.5_ represents the value of depending on C when the coverage is smaller than 0.5. Note that *β* does not have a subscript because it does not change across the whoel range of coverage values so it represents the average value of the exponent for coverages from 0-1. Given that the prefactor is approximately constant for *C >* 0.5, we fitted Eq. 7 for samples with coverages *>* 0.5 and estimated the following parameters values; *α*_*C>*0.5_ = 10^0.97^, *β* = 0.35 [19]. Thus for coverages 0.5 the prefactor and exponent are constant but for coverages *<* 0.5 the prefactor increases linearly with C,

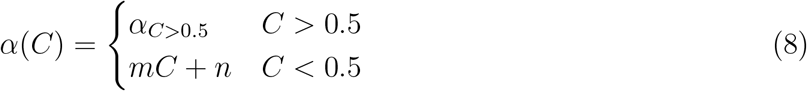

where *m* = 12.95, *n* = −1.17, which were estimated by allowing the exponent to vary. (The parameters with a fixed exponent were similar, *m* = 17.17, *n* = −0.86). Based on the number of protein types that a given total number of protein molecules should have, it is possible to write an equation to predict the number of expressed proteins. The equation to calculate a maximum (for *C >* 0.5) number of proteins, *D*^*′*^(*N, λ*(*α*), *α*(*C*)), taking into account the coverage is,

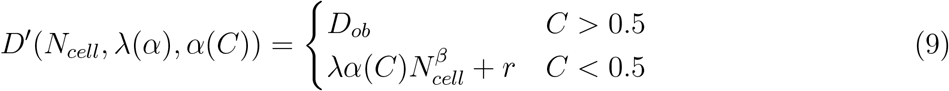

where *D*_*ob*_ is the observed value of a point in the relationship according to eq. 9 and *D*(*N, α*(*C <* 0.5)) is the predicted value by Eq. 9,

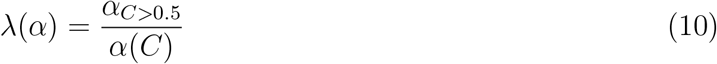

The parameters for this equation are 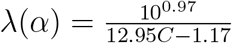. The residuals (r) are defined as,

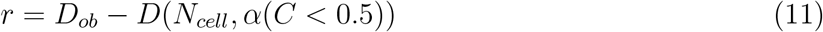

Eq. 9 allows scale data on the number of proteins per cell of samples with coverages lower than 0.5, assuming that below this point, the number of proteins is limited by experimental limitations, such as sample amount, which affects number of cells in the sample.

## Algorithm

The algorithm is summarized in Table 2. The input is genome size (Mbp) and the vector of protein abundances in the sample. It consists of basically two steps; 1) From genome size, and based on scaling relationships of Eqs. 1-6, it calculates the total proteins in the cell, the abundance of each protein per cell, the number of protein-coding genes, and the coverage. 2) Then, there is a conditional where if the coverage is greater or lower than 0.5, the number of protein types or maximum possible observed protein types (for a coverage greater than 0.5) is calculated. Finally, the output is the total number of proteins, the vector of individual protein abundances, and the number of protein types for a coverage greater than 0.5. We implemented the algorithm as a function in the R language. We provide the R code and a couple of examples that show how to use it the Supplementary Material.

### Validation

To assess how different the estimations for proteome properties are compared to other methods, we applied our method to *E. coli K-12* and compared the estimations made by Milo (2013) and the recent Single-Cell (Vegvari et al. 2023) approach. Milo et al. 2013 estimated a total protein of 3*×*10^6^ and Vegvari et al. 2023 a total of 3.88*×*10^5^, compared to our estimation of 8.13*×*10^6^.

## Discussion

Here we have used scaling relationships between genome size, protein volume, cell volume, and protein types, among others, to develop a simple algorithm to compute per-cell properties of proteomes; including the total number of protein molecules per cell, the abundance per cell of each individual protein, and the number of protein types.

Some of the applications of this method could predict the properties of natural and artificial cells. For the design of artificial cells, for example, it easily predicts the total number of proteins or different types. This method can also be easily extended for unicellular eukaryotes, or even for multicellular ones, given that there are scaling relationships for these cells [20]. There are some considerations. For the same species, cell volume changes with nutrient concentration, temperature, etc, and most of the data on which these relationships are based correspond to compilations of the literature based on different conditions. Despite this, the scaling relationships are valid across a wide range of species and the variance could correspond to that intraspecific variation due to stochasticity and environmental variation. Also, in this approach, we predict how many different proteins in a cell there should be but not the relative abundance of unobserved proteins. In synthesis, we developed a method, based on cell scaling relationships, to calculate the total number of proteins, the number of different proteins, and the abundance of each protein per cell. The code is written in the R environment and be easily implemented with just five line codes as detailed above. This method has applications in the design of artificial cells as a predictive framework to calculate, for example, the concentrations of specific proteins of protein functions, and can be extended to eukaryotes. As final comments, our study is a good example of the application of scaling laws to make applied predictions. However, despite each of these relationships being well-established empirically, still is needed to develop a theory from first principles to understand the origin of most of these relationships.

## Acknowledgments

NSF projects: ‘Building and Modeling Synthetic Bacterial Cells’ (1840301), and ‘Towards a unified theory of regulatory functions and networks across biological and social systems’ (2133863).

## Code and data availability

The code and data will be available at: https://github.com/jose-ignacio-arroyo/proteomic-data-scaling

