## Supplementary Material for "An algorithm for predicting per-cell proteomic properties"

### Supplementary Material
### An algorithm for per-cell proteomic data scaling

José Ignacio Arroyo (*), Chris Kempes, and Geoffrey West
Santa Fe Institute, NM, USA
(*)

#### Supplementary Figures

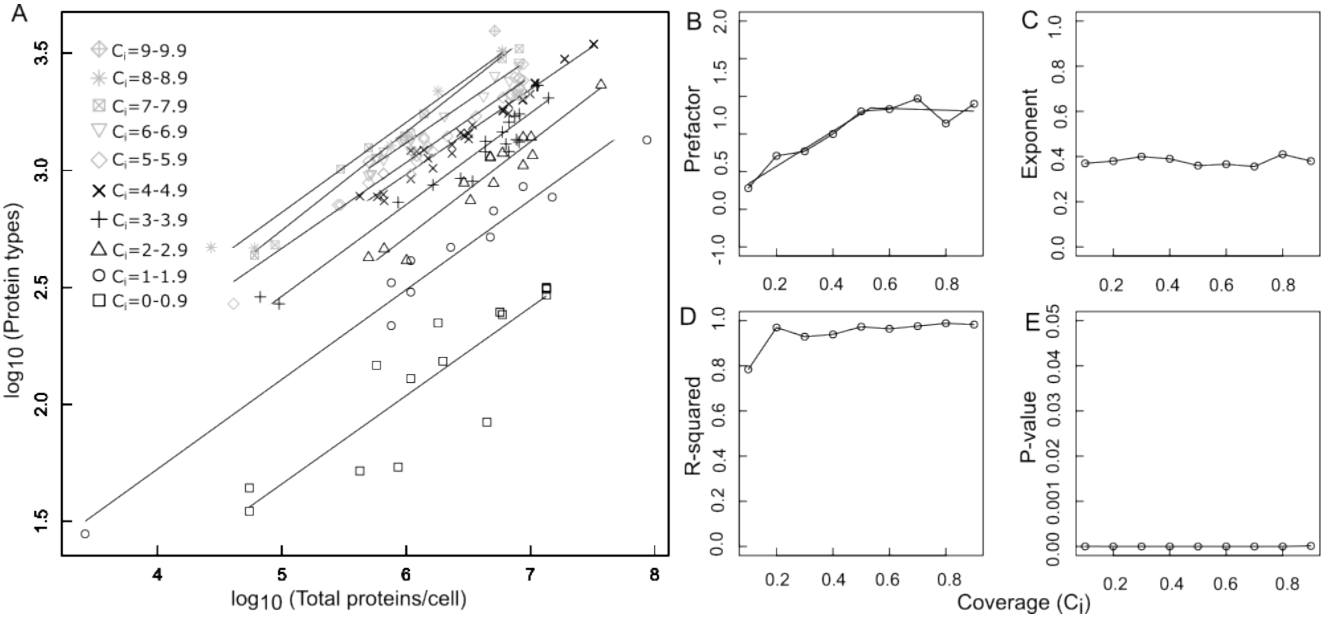


Figure 1: a) Relationship between protein types and total proteins, $D = \alpha N^{\beta}$, where $\alpha$ is the prefactor and $\beta$ the exponent. The points in gray have coverage above 0.5, and each of the different symbols denotes a different coverage interval. It can be seen that for data above a coverage of 0.5, there is a very well-defined relationship. Relationship between b) $a$ and coverage. The fitted line corresponds to a segmented regression that estimated a breakpoint at a coverage of 0.53. c) exponent and coverage, d) r-squared and coverage, and e) p-value and coverage.

#### Supplementary Methods

**Implementation in R.** We wrote an algorithm (in the R language) that takes as an input the vector of protein abundances or counts and the genome size, and outputs the total protein abundance per cell, the abundance of each cell and the number of different expressed proteins. The key calculations rely on well-established empirical cell scaling relationships. It should be noted that the scaling relationships described here allow for other inputs beyond genome size. For example, if cell volume is directly measured it could be used instead, and the key technique is to use the scaling relationships to estimate whichever features are unmeasured. Below, we explain each of the steps of the algorithm and how to use it in the R environment.

The algorithm was implemented as a function "pds" that we wrote in the R language available in the file "pds.R" which is downloadable from `https://github.com/jose-ignacio-arroyo/proteomic-data-scaling` or the Supplementary Material of this article. We write the example code in `Courier` and the comments in normal characters.

To run the code in an R terminal first upload the code of the algorithm,

```
source("pds.R")
```

The function needs two inputs, the genome size (Mb) and the vector of protein abundances. These data can be obtained from the NCBI site. Upload the vector of the protein abundances or counts, using for example the function "read.table" and defining the vector,

```
df=read.table("exampleE.coliK12.txt", h=T)

vecpro=df[,1]
```

Define the genome size value. For example for *E. coli* is 5.12,

```
mb=5.12
```

Now we just apply the function "pds" (which inputs are "vecpro" and "mb") are and we can define a name for the ouput. let's just call it "pds.output",

```
pds.output= pds(mb, vecpro)
```

The output, "pds.output" (not displayed here), is a list containing three elements; the predicted total number of proteins per cell, the predicted number of different proteins per cell, and a vector of scaled protein abundances per cell.
